# Microtubule poleward flux as a target for modifying chromosome segregation errors

**DOI:** 10.1101/2024.03.11.584498

**Authors:** Patrik Risteski, Jelena Martinčić, Erna Tintor, Mihaela Jagrić, Iva M. Tolić

**Affiliations:** Division of Molecular Biology, Ruđer Bošković Institute, Bijenička cesta 54, 10000 Zagreb, Croatia

## Abstract

Cancer cells often display errors in chromosome segregation, some of which result from improper chromosome alignment at the spindle midplane. Chromosome alignment is facilitated by different rates of microtubule poleward flux between sister kinetochore fibers. However, the role of the poleward flux in supporting mitotic fidelity remains unknown. Here, we introduce the hypothesis that the finely tuned poleward flux safeguards against lagging chromosomes and micronuclei at mitotic exit by promoting chromosome alignment in metaphase. We used human untransformed RPE-1 cells depleted of KIF18A/kinesin-8 as a system with reduced mitotic fidelity, which we rescued by three mechanistically independent treatments, comprising low-dose taxol or co-depletion of the spindle proteins HAUS8 or NuMA. The rescue of mitotic errors was due to shortening of the excessively long overlaps of antiparallel microtubules, serving as a platform for motor proteins that drive the flux, which in turn slowed down the overly fast flux and improved chromosome alignment. In contrast to the prevailing view, the rescue was not accompanied by reduction of overall microtubules growth rates. Speckle microscopy revealed that the improved chromosome alignment in the rescue treatments was associated with slower growth and flux of kinetochore microtubules. In a similar manner, a low-dose taxol treatment rescued mitotic errors in a high-grade serous ovarian carcinoma cell line OVKATE. Collectively, our results highlight the potential of targeting microtubule poleward flux to modify chromosome instability and provide insight into the mechanism through which low doses of taxol rescue mitotic errors in cancer cells.

## Introduction

To achieve proper inheritance of the duplicated genome, the cell aligns chromosomes at the spindle equator before dividing them between the daughter cells. Chromosome alignment at the metaphase plate promotes synchronous poleward movement of chromosomes in anaphase, and thus their faithful segregation (Maiato et al., 2017). A precisely formed metaphase plate is critical for the maintenance of genome stability because misaligned chromosomes may lag behind separating chromosome masses or remain near the pole throughout mitosis to form micronuclei (Fonseca et al., 2019; Gomes et al., 2022; Marquis et al., 2021; Quinton et al., 2021; Tucker et al., 2023), which are prone to acquire DNA damage and undergo chromosomal rearrangements, including chromothripsis (Cortés-Ciriano et al., 2020; Liu et al., 2018; Zhang et al., 2015). Therefore, chromosome alignment defects are considered to be one of the origins of chromosomal instability (CIN) in cancer cells (Gordon et al., 2012; Thompson et al., 2010) and are implicated in the development of oncogene-independent selective advantage (Rowald et al., 2016).

Essential to chromosome alignment on the spindle are microtubules and microtubule-associated proteins, which jointly exert forces on kinetochores and chromosome arms to maintain their position at the spindle equator (Risteski et al., 2021). Regulation of the microtubule growth rates and stability is crucial in the context of chromosome missegregation because excessively fast microtubule growth is a feature of cancer cells that exhibit CIN, and taxol treatment can rescue chromosome missegregation by slowing down microtubule growth (Bakhoum et al., 2009; Ertych et al., 2014; Tamura et al., 2020). Microtubule dynamics is modulated by a variety of proteins, with the most prominent player being the motor protein KIF18A/kinesin-8. This protein suppresses polymerization of microtubule plus ends (Du et al., 2010; Stumpff et al., 2012), thereby limiting kinetochore movements (Stumpff et al., 2008). Recent studies identified aneuploid cancer cells to be particularly sensitive to the depletion or inhibition of KIF18A (Cohen-Sharir et al., 2021; Marquis et al., 2021; Quinton et al., 2021). Interestingly, in CIN tumor cells that exhibit elevated spindle microtubule dynamics, KIF18A depletion can be exploited to reduce their proliferation, by increasing the already elevated microtubule dynamics, which leads to increased chromosome misalignment and therefore CIN and cell death (Quinton et al., 2021). This is in agreement with the sensitization of tumor cells by elevating the frequency of chromosome missegregation (Kops et al., 2004; Janssen et al., 2009), as high rates of missegregation suppress tumor progression (Silk et al., 2013; Zasadil et al., 2016). Thus, microtubule dynamics and, consequently, chromosome alignment have a direct effect on CIN levels in cancers.

Apart from its role via suppressing microtubule plus-end dynamics, KIF18A is implicated in chromosome alignment on the spindle via overlap length-dependent forces (Jagrić et al., 2021; Risteski et al., 2022). In this model, kinetochore fibers (k-fibers), which are attached end-on to sister kinetochores, are laterally linked by a bundle of non-kinetochore microtubules called the bridging fiber (Kajtez et al., 2016; Pavin and Tolić, 2021; Tolić, 2018). Motor proteins within the overlaps between bridging and k-fibers slide the fibers apart (Risteski et al., 2022), thereby generating their movement towards the poles, known as poleward flux (Forer, 1965; Hamaguchi et al., 1987; Hiramoto and Izutsu, 1977; Mitchison, 1989). The flux of longer k-fibers is faster due to their longer overlaps with bridging fibers, resulting in kinetochore movement towards the spindle midplane (**Fig. 1A**, scheme). In contrast, when the overlaps between the bridging and k-fibers are excessively long, as after KIF18A depletion, the coupling between the fibers becomes overly strong and sister k-fibers undergo equally fast flux, leading to inefficient chromosome alignment (Risteski et al., 2022). Thus, chromosome misalignment can be a consequence of altered microtubule poleward flux.

**Fig. 1.**
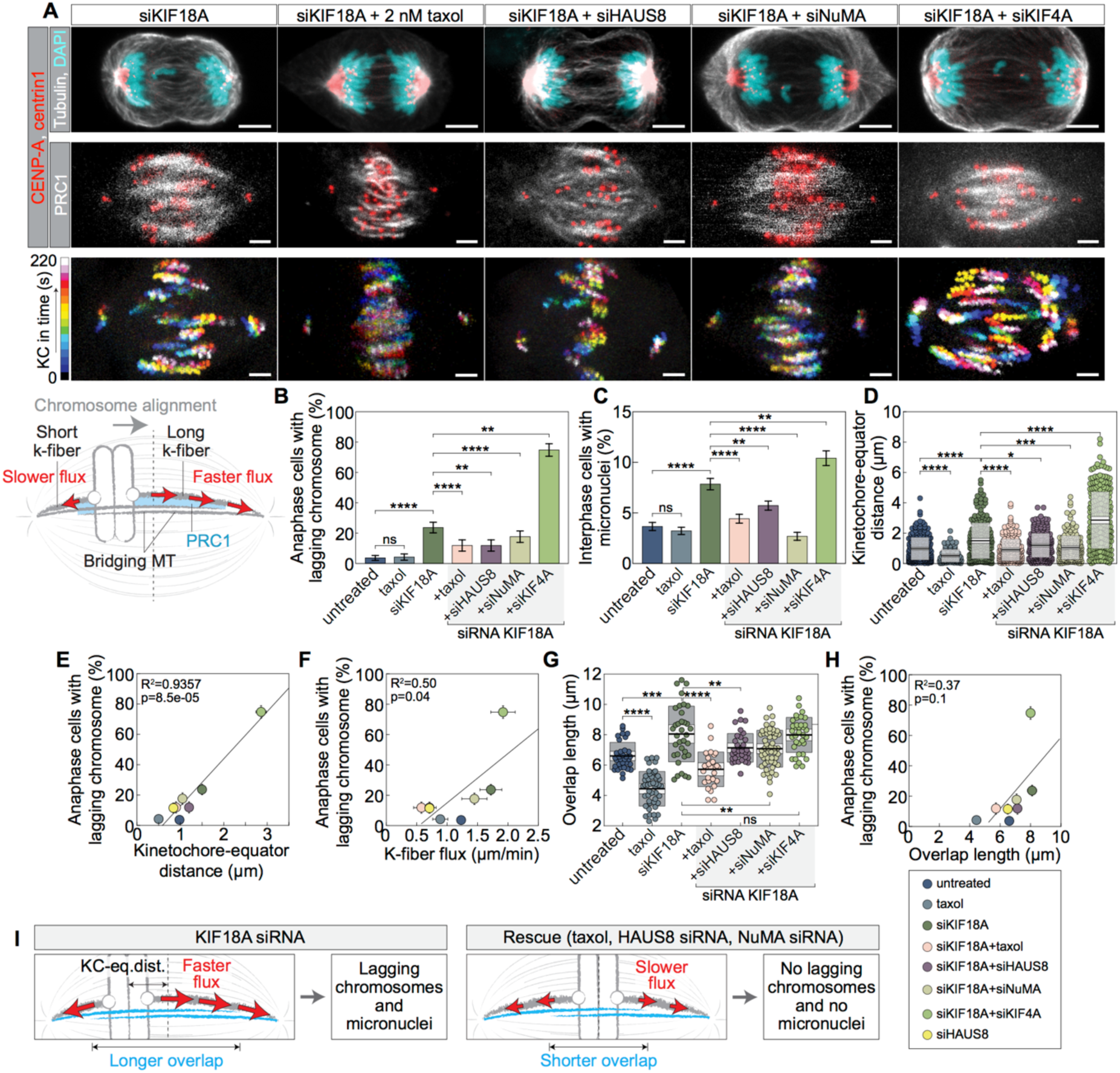
Lagging chromosomes and micronuclei in KIF18A-depleted cells can be rescued by decelerating microtubule poleward flux. **A,** RPE-1 cells stably expressing CENP-A-GFP and centrin1-GFP (top, middle: red; bottom: temporal color projection) for each respective siRNA treatment. The cells are immuno-stained for tubulin-AF-594 (top: grey) or PRC1-AF-594 (middle: grey) and dyed with DAPI (top: cyan). Scale bars: 5 µm (top), 2 µm (middle, bottom). Scheme depicts the flux-driven centering model. When a kinetochore pair is off-centered, the longer k-fiber exhibits a more extended overlap with the bridging fiber, strengthening the coupling and causing a faster flux compared to the shorter sister k-fiber. Consequently, the kinetochore pair moves in the direction of the longer k-fiber, i.e., towards the spindle midplane. **B,** Percentage of anaphase cells with lagging chromosomes, color-coded for treatments. Number of analyzed cells for treatments from left to right: 138, 95, 148, 76, 76, 102, 83, 96. **C,** Percentage of interphase cells with micronuclei, color-coded for treatments. Number of analyzed cells for treatments from left to right: 2215, 2421, 2269, 2215, 2675, 1676, 1768, 2455. **D,** Distance between sister kinetochore pair midpoints and the spindle equator, color-coded for treatments. Number of distances measured for sister kinetochore pairs from the equator for treatments from left to right: 258, 97, 198, 190, 242, 191, 235, 137. **E,** Percentage of anaphase cells with lagging chromosomes plotted against the kinetochore-equator distance, color-coded for treatments. **F,** Percentage of anaphase cells with lagging chromosomes plotted against the k-fiber flux, color-coded for treatments. **G,** PRC1-labeled fiber overlap length, color-coded for treatments. Number of PRC1-labelled bundles measured for treatments from left to right: 33, 45, 35, 27, 35, 53, 35, 30. **H,** Percentage of anaphase cells with lagging chromosomes plotted against the PRC1-labeled overlap length, color-coded for treatments. **I,** Schematic showing that increased microtubule poleward flux leads to increased chromosome misalignment on the spindle, which in turn causes higher frequencies of lagging chromosomes in anaphase and micronuclei in the next cell cycle. Legend is at the bottom right corner. Maximum intensity z-projection (A; top, middle). Kinetochore-equator distances, overlap lengths, and k-fiber flux for untreated, siKIF18A, siKIF18A+siHAUS8, siKIF18A+siKIF4A, and siHAUS8 were replotted from (Risteski et al., 2022). Statistical tests: two-proportions z-test (B,C), Mann-Whitney test (D), t-test (G). p=p-value, R^2^=coefficient of determination. p(ns)>0.05, p(*)=0.01–0.05, p(**)=0.01–0.001, p(***)=0.001–0.0001, p(****)<0.0001.

Besides KIF18A, various other microtubule-associated proteins can modify the microtubule poleward flux, including HAUS6/8 (Almeida et al., 2022; Risteski et al., 2022; Štimac et al., 2022), NuMA (Risteski et al., 2022; Steblyanko et al., 2020; Sun et al., 2021), and KIF4A (Risteski et al., 2022; Steblyanko et al., 2020). HAUS8 is a subunit of the augmin complex, which promotes microtubule nucleation along the walls of other microtubules (Goshima et al., 2008; Kamasaki et al., 2013; Wu et al., 2009), particularly the nucleation of bridging fibers (Štimac et al., 2022). NuMA serves as a passive crosslinker for parallel microtubules, working in tandem with dynein to tether microtubules towards the spindle poles (Hueschen et al., 2017; Merdes et al., 1996). The coupling between bridging fibers and k-fibers through NuMA promotes the force transmission to k-fibers and therefore k-fiber flux (Risteski et al., 2022; Steblyanko et al., 2020). The forces driving flux are also affected by KIF4A, a plus-end directed motor that inhibits the length and growth rate of non-kinetochore microtubules (Jagrić et al., 2021; Stumpff et al., 2012; Wandke et al., 2012), thereby shortening bridging fiber overlaps and in turn slowing down k-fiber flux (Risteski et al., 2022). In addition to regulating the flux, these proteins have been implicated in chromosome alignment and the occurrence of chromosome segregation errors (Gomes et al., 2022; Maiato et al., 2017; Risteski et al., 2022). However, the role of microtubule poleward flux in safeguarding mitotic fidelity is unknown.

Here, we probe the idea that microtubule poleward flux promotes chromosome inheritance fidelity by helping chromosome alignment, and explore the potential of targeting flux to reduce mitotic errors. In non-transformed cells, lagging chromosomes and micronuclei induced by KIF18A depletion were partially rescued by a low dose of taxol or co-depletion of HAUS8 or NuMA. The treatments that rescued mitotic errors in KIF18A-depleted cells did so by shortening the antiparallel overlaps and slowing down the flux and growth of kinetochore microtubules, thereby improving chromosome alignment, rather than by slowing down microtubule growth in general. Reducing overlaps and flux rates restored chromosome alignment and improved segregation fidelity in an ovarian cancer cell line. These findings highlight the potential to modify CIN in cancer cells by targeting microtubule poleward flux.

## Results

### Lagging chromosomes and micronuclei caused by chromosome misalignment can be rescued by slowing down microtubule poleward flux

This study posits the following hypothesis: faster poleward flux of spindle microtubules leads to chromosome misalignment in metaphase, which in turn results in lagging chromosomes during anaphase and micronuclei in the following interphase. The primary implication of this hypothesis is that alterations to the microtubule poleward flux have the potential to rescue these mitotic errors.

To explore our hypothesis, we used human non-transformed hTERT-immortalized retinal pigment epithelium cells (RPE-1), which we depleted of KIF18A because this depletion leads to chromosome misalignment (Garcia et al., 2002; Mayr et al., 2007; Stumpff et al., 2012, 2008; West et al., 2002; Zhu et al., 2005), occurrence of lagging chromosomes and micronuclei (Fonseca et al., 2019; Garcia et al., 2002; Stumpff et al., 2008; West et al., 2002), and excessively fast poleward flux of k-fibers (Risteski et al., 2022). To test whether lagging chromosomes and micronuclei in this system can be rescued by additional treatments, we used three mechanistically independent approaches: (i) we treated KIF18A-depleted cells with a low dose of paclitaxel (taxol), a treatment that partially rescues mitotic errors in cancer cells with increased microtubule dynamics (Crowley et al., 2022; Tamura et al., 2020); (ii) we co-depleted HAUS8, a subunit of the augmin complex, because this co-depletion rescues chromosome misalignment of KIF18A-depleted cells (Risteski et al., 2022); (iii) we co-depleted NuMA, which regulates the poleward flux of k-fibers (Risteski et al., 2022) (**Fig. 1A**; Supplementary Fig. S1A–S1D). Remarkably, all three treatments alleviated the increased incidence of both lagging chromosomes and micronuclei induced by the depletion of KIF18A (**Fig. 1B and C**). Moreover, these treatments improved the alignment of chromosomes at the metaphase plate in KIF18A-depleted cells (**Fig. 1A and 1D**; Supplementary Fig. S1D; Supplementary Movie 1). As a positive control, we used co-depletion of kinesin-4/KIF4A and KIF18A, which increases chromosome misalignment in comparison with KIF18A depletion (Risteski et al., 2022), and found that this co-depletion further increased the occurrence of lagging chromosomes and micronuclei (**Fig. 1A–D**). Across all conditions, the degree of chromosome misalignment correlated with the incidence of lagging chromosomes and with the occurrence of interphase micronuclei (**Fig. 1E**; Supplementary Fig. S1E). The latter two phenomena were also correlated (Supplementary Fig. S1F), in agreement with previous studies linking chromosome segregation errors and micronuclei (Gomes et al., 2022; Thompson and Compton, 2011). Overall, our findings suggest that the mitotic errors in KIF18A-depleted cells can be mitigated by improving chromosome alignment on the metaphase spindle.

Given the crucial role of microtubule poleward flux in promoting chromosome alignment on the spindle (Risteski et al., 2022, 2021) and the anaphase synchronization of chromosome movements in anaphase (Matos et al., 2009), we explored whether the observed changes in chromosome missegregation can be explained by the changes in microtubule poleward flux rates. By using the recently developed dye-based fluorescence speckle microscopy (Risteski, 2023; Risteski et al., 2022), we measured the poleward movement of individual speckles on the microtubule lattice. The movement of each speckle represents the poleward flux of the microtubule carrying the speckle, allowing for separate measurements of kinetochore microtubule flux and bridging microtubule flux (Supplementary Fig. S1G and H). Across all conditions, the relationships between the microtubule flux parameters were in agreement with the flux-driven kinetochore centering model (Supplementary Fig. S1I and J) (Risteski et al., 2022). Interestingly, the k-fiber flux rates correlated with the incidence of lagging chromosomes (**Fig. 1F**; Supplementary Fig. S1K), suggesting that flux perturbations affect chromosome inheritance fidelity.

As the increased microtubule poleward flux in KIF18A-depleted cells was proposed to be a result of extended overlaps between the antiparallel microtubules and thus higher sliding forces (Jagrić et al., 2021; Risteski et al., 2022), we set out to determine the size of these overlaps in treatments that rescued mitotic errors. We found that the antiparallel overlap length, measured as the length of stripes decorated by the crosslinker of antiparallel microtubules PRC1 (Kajtez et al., 2016; Polak et al., 2017), was reduced in KIF18A-depleted cells after the low-dose taxol treatment or after co-depletion with HAUS8 or NuMA (**Fig. 1A and G**; Supplementary Fig. S1D). These results indicate that the extended antiparallel overlaps play a role in generating mitotic errors in KIF18A-depleted cells. Across all conditions, longer overlaps tended to coincide with faster poleward flux, worse chromosome alignment, and a greater prevalence of lagging chromosomes and micronuclei, though the correlation was statistically significant only for chromosome alignment (**Fig. 1H**; Supplementary Fig. S1L–N).

Taken together, our results suggest that overlap length-dependent microtubule poleward flux safeguards mitotic fidelity by promoting chromosome alignment. It is important to acknowledge that, in principle, each specific treatment applied here could have influenced mitotic fidelity independently of the overlap length and poleward flux. Nonetheless, the fact that three mechanistically independent approaches rescued both the flux-dependent chromosome alignment and segregation errors supports the causal connection between microtubule poleward flux and mitotic fidelity (**Fig. 1I**).

### Altered overall growth rates of microtubules cannot explain the changes in chromosome alignment and missegregation

Aberrant microtubule dynamics and stability are features of cancer cells that lead to chromosomal instability, and taxol treatment can rescue chromosome missegregation by reducing excessive microtubule growth velocities (Bakhoum et al., 2009; Ertych et al., 2014; Tamura et al., 2020). However, it is unclear how the treatments that affect microtubule dynamics, including taxol and our co-depletions, rescue mitotic errors. There are two main possibilities: First, the primary mechanism of action may involve a reduction in the growth rate of all microtubules. In this scenario, treatments with a more pronounced slowdown of overall microtubule growth should more efficiently rescue mitotic errors. Second, these treatments may influence mitotic fidelity by decreasing the plus-end growth rate of kinetochore microtubules. This rate depends not only on intrinsic microtubule growth but also on the forces exerted by the kinetochore onto the microtubule plus ends. These forces, in turn, are dependent on the microtubule poleward flux. In this scenario, treatments with a more pronounced slowdown of the poleward flux and of the kinetochore microtubule growth velocity should more efficiently rescue the errors.

To distinguish between these possibilities, we first measured growth velocities of astral microtubules on metaphase spindles, as a readout of the general microtubule growth, in each condition. For this purpose, we tracked spots of GFP-labeled EB3, which marks the plus ends of growing microtubules (**Fig. 2A**; Supplementary Fig. S2A; Supplementary Movie 2) (Stepanova et al., 2003). Taxol decreased EB3 spot velocities in wild-type cells (Ertych et al., 2014), and KIF18A depletion had no effect (**Fig. 2B**) (Stumpff et al., 2012). Interestingly, the treatments that rescued mitotic errors in KIF18A-depleted cells either did not change (taxol) or increased the EB3 spot velocities (HAUS8 or NuMA co-depletion) (**Fig. 2B**). Across conditions, the EB3 spot velocity was not predictive of the efficiency of chromosome alignment (**Fig. 2C**). These results are not consistent with the hypothesis that a general perturbation of the microtubule plus-end dynamics underlies the improved chromosome alignment and mitotic fidelity following taxol treatment, as well as after co-depletion of HAUS8 or NuMA, in KIF18A-depleted cells.

**Fig. 2.**
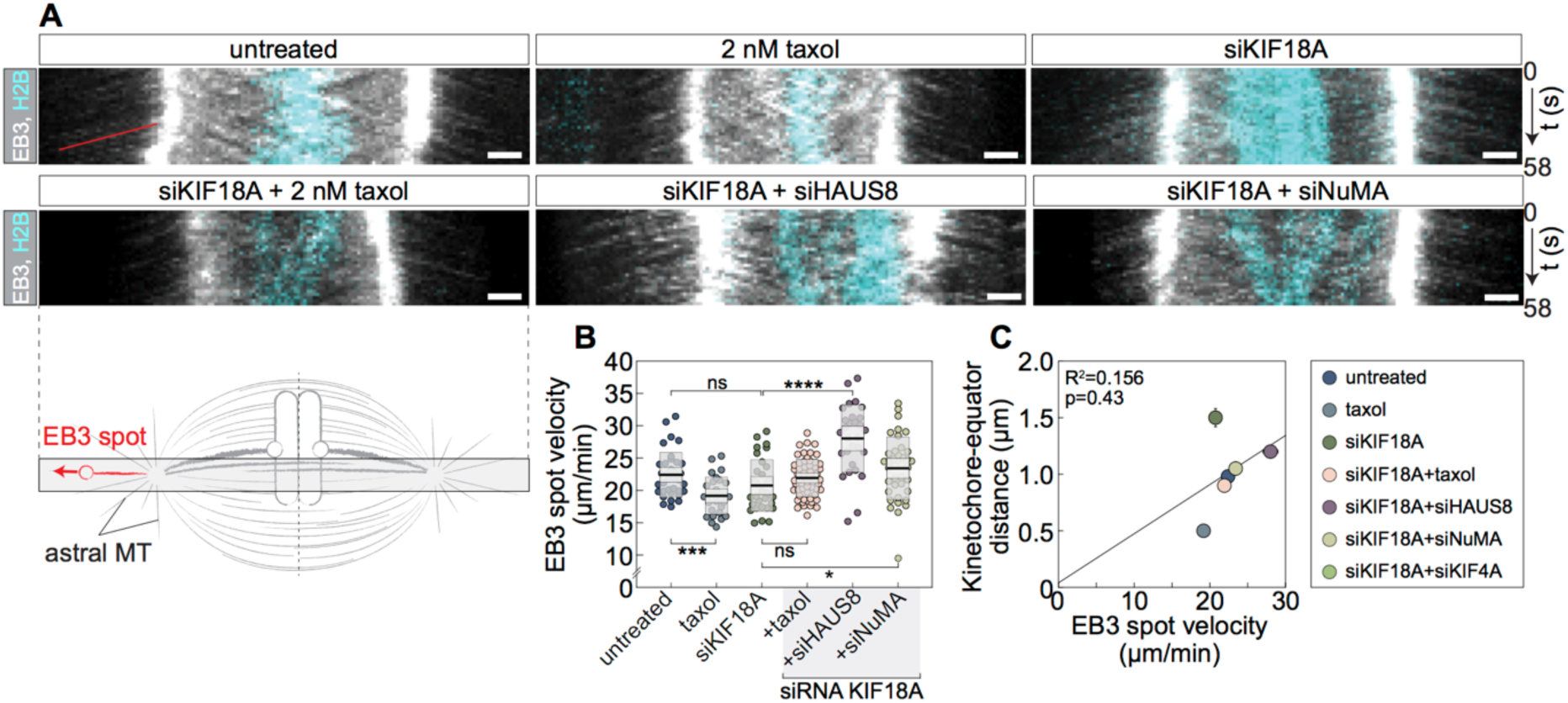
Treatments that reduce microtubule poleward flux do not reduce astral microtubule plus-end dynamics. **A,** Kymographs (temporal projections) of RPE-1 cells with stable expression of EB3-GFP (grey) and H2B-mCherry (cyan), per corresponding treatment. Scale bars: 2 µm. The red line marks a representative EB3 trace over time. The scheme indicates the position at which the kymographs were created. **B,** EB3 spot velocity, measured on astral microtubules of metaphase spindles. Number of EB3 spots measured for treatments from left to right: 35, 28, 31, 57, 29, 38. **C,** Kinetochore-equator distance plotted against the EB3 spot velocity, color-coded for treatments. Legend is at the bottom right corner. Kinetochore-equator distances for untreated, siKIF18A, siKIF18A+siHAUS8, and siKIF18A+siKIF4A were replotted from (Risteski et al., 2022). Statistical tests: t-test (B). p-value as in Fig. 1.

### Altered kinetochore microtubule growth and flux can explain the changes in chromosome alignment and missegregation

To explore the hypothesis that taxol and HAUS8 or NuMA co-depletions influence mitotic fidelity by decreasing the plus-end growth rate of k-fibers, we used speckle microscopy to measure growth or shrinkage rates of kinetochore microtubules at their plus end, i.e., at the kinetochore (**Fig. 3A**). Given that pairs of sister kinetochores oscillate around the spindle midplane during metaphase (Amaro et al., 2010; Armond et al., 2015; Jaqaman et al., 2010; Stumpff et al., 2008), we assessed microtubule dynamics separately at the leading and the trailing kinetochore, where the leading kinetochore refers to the one moving toward its associated pole, while the trailing kinetochore is the one moving away from its linked pole. We measured the velocity with which a speckle, fixed on the lattice of a kinetochore microtubule, moves with respect to its kinetochore (**Fig. 3A**). If the speckle moves away from the kinetochore, we interpret this as growth of the kinetochore microtubule. Conversely, if the speckle moves towards the kinetochore, we consider it as shrinkage of the kinetochore microtubule.

**Fig. 3.**
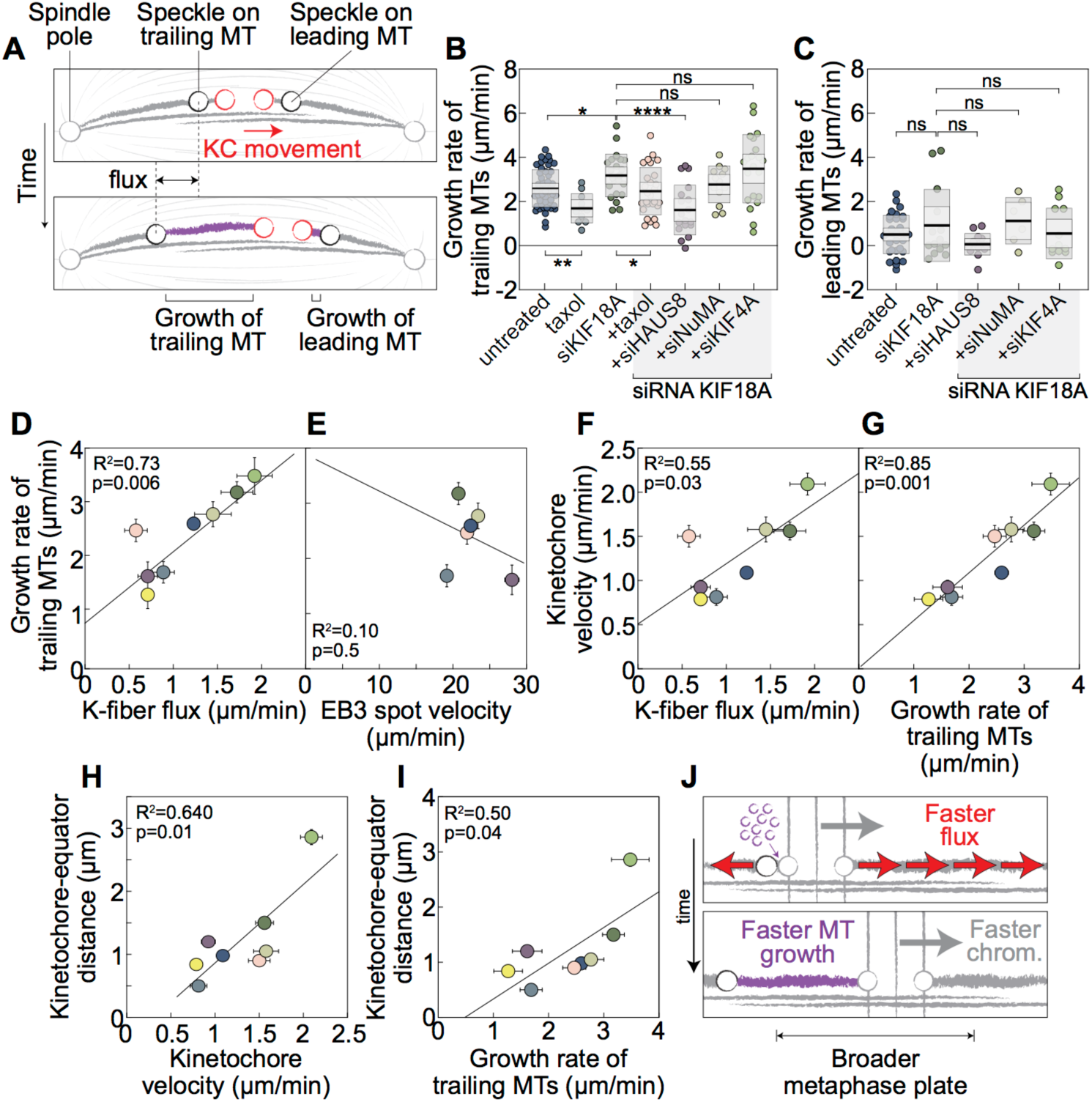
Altered microtubule poleward flux affects the metaphase plate width and kinetochore velocities via its effect on k-fiber growth rates. **A,** Schematic depiction illustrating the differentiation between leading and trailing kinetochores, along with their respective associated k-fibers and bridging fibers. **B,** Growth rate of microtubules measured as speckle-to-kinetochore velocity on k-fiber associated with trailing kinetochore. Number of analyzed k-fiber speckles associated with trailing kinetochores for treatments from left to right: 48, 11, 24, 27, 18, 13, 21, 14. **C,** Growth rate of microtubules measured as speckle-to-kinetochore velocity on k-fiber associated with leading kinetochore. Number of analyzed k-fiber speckles associated with leading kinetochores for treatments from left to right: 32, 14, 14, 6, 12, 16. D, Growth rate of microtubules associated with trailing kinetochore plotted against the k-fiber flux rate, color-coded for treatments. **E,** Growth rate of microtubules associated with trailing kinetochore plotted against the EB3 spot velocity, color-coded for treatments. **F,** Kinetochore velocity plotted against the k-fiber flux rate, color-coded for treatments. **G,** Kinetochore velocity plotted against the growth rate of microtubules associated with trailing kinetochore, color-coded for treatments. H, Kinetochore-equator distance plotted against the kinetochore velocity, color-coded for treatments. **I,** Kinetochore-equator distance plotted against the growth rate of microtubules associated with trailing kinetochore, color-coded for treatments. J, Schematic showing that microtubule poleward flux leads to microtubule growth at the plus-ends of k-fibers associated with trailing kinetochores and in turn kinetochore movement within the metaphase plate. Note that kinetochore-equator distances and k-fiber flux rates for untreated, siKIF18A, siKIF18A+siHAUS8, siKIF18A+siKIF4A, and siHAUS8 were replotted from (Risteski et al., 2022). Statistical tests: t-test (B,C). p-value as in Fig. 1.

We found that the trailing kinetochore microtubules grew at the plus end under all examined conditions (**Fig. 3B**). Taxol treatment reduced their growth rate, whereas KIF18A depletion increased it. Furthermore, taxol treatment and HAUS8 co-depletion decreased the excessively high kinetochore microtubule growth rate of KIF18A-depleted cells. Co-depletion of NuMA caused slightly slower growth, though the effect was not significant (**Fig. 3B**).

In contrast to the trailing kinetochore microtubules, their leading counterparts showed slow growth at the plus end, which was similar across all treatments (**Fig. 3C**). This result suggests that the leading kinetochore microtubules are typically in a state of slow growth or pausing at their plus end, and that the used treatments did not affect their plus-end dynamics. It is worth noting that the results for speckles on the leading kinetochore microtubules might be biased towards higher microtubule growth velocities because speckles that disappeared within the first 30 seconds after appearance, due to depolymerization of the leading microtubule plus end, were too short-lived to be analyzed. However, such disappearance events were rare, suggesting that the bias was not substantial. As the measurements of the trailing microtubule growth velocities were unbiased, because these speckles moved away from their kinetochore, and more abundant, we use them as representative of kinetochore microtubule plus-end growth in the subsequent analysis.

Across conditions, kinetochore microtubule plus-end growth velocity strongly correlated with the k-fiber flux velocity (**Fig. 3D**), but did not correlate with EB3 spot velocity (**Fig. 3E)**. These results indicate that the changes in the kinetochore microtubule plus-end growth are mainly determined by the altered forces acting at and near the kinetochore, rather than by the altered intrinsic growth of free plus ends in each condition.

The velocity of kinetochore movements correlated with the k-fiber flux and the kinetochore microtubule plus-end growth velocity across treatments (**Fig. 3F and G**; Supplementary Fig. S3A). Interestingly, we found that treatments with increased kinetochore velocities, as well as those with increased kinetochore microtubule plus-end growth velocities, tend to have worse chromosome alignment (**Fig. 3H and I**). This is in contrast with the absence of a correlation between EB3 spot velocity and the efficiency of chromosome alignment (**Fig. 2C**). We propose that a faster poleward flux is accompanied by a faster growth of the kinetochore microtubule plus end, allowing for faster kinetochore movements during their oscillations around the spindle midplane, ultimately resulting in a broader metaphase plate (**Fig. 3J**). Thus, altered poleward flux of kinetochore microtubules due to changes in the antiparallel overlaps, rather than a general perturbation of the microtubule plus-end dynamics, can explain the improved chromosome alignment and mitotic fidelity following taxol treatment, as well as after co-depletion of HAUS8 or NuMA, in KIF18A-depleted cells.

### Chromosome missegregation rates in cancer cells can be reduced by altering microtubule poleward flux

To investigate the relevance of our findings on the protective role of microtubule poleward flux in maintaining mitotic fidelity for cancer cells, we set out to rescue mitotic errors in specific cancer cell lines. To find promising candidate lines, we performed a mini screen where we roughly assessed the presence of wider metaphase plates and longer spindles, using the latter as an indicator of extended antiparallel overlaps (Supplementary Fig. S4). We tested the following lines: rhabdomyosarcoma (RD), colorectal adenocarcinoma (DLD-1), lung adenocarcinoma (A549), and ovarian carcinoma (OVKATE and OVSAHO) (Supplementary Fig. S4A). OVKATE was identified as the most promising candidate, prompting us to select it along with its counterpart, OVSAHO, both serving as models of high-grade serous ovarian carcinoma (Barnes et al., 2021; Domcke et al., 2013).

We aimed to assess whether targeting microtubule poleward flux could reduce chromosome segregation errors in OVKATE and OVSAHO cells. We found that a low-dose taxol treatment decreased the occurrence of lagging chromosomes in OVKATE cells, whereas in OVSAHO cells this decrease was smaller and not significant (**Fig. 4A**; Supplementary Fig. S4B and C; our result in OVSAHO differs from a previous study (Tamura et al., 2020), which found a significant reduction of anaphase errors). In line with fewer lagging chromosomes, OVKATE cells also had fewer micronuclei after taxol treatment (**Fig. 4B**). In contrast to OVKATE, the number of micronuclei in OVSAHO cells increased after taxol treatment (**Fig. 4B**), possibly due to the stabilization of BRCA2-driven uncongressed chromosomes and premature chromosome segregation in this cell line (Choi et al., 2012; Ehlén et al., 2020; Tamura et al., 2020), which can cause the formation of micronuclei (Choi et al., 2012; Ehlén et al., 2020).

**Figure 4.**
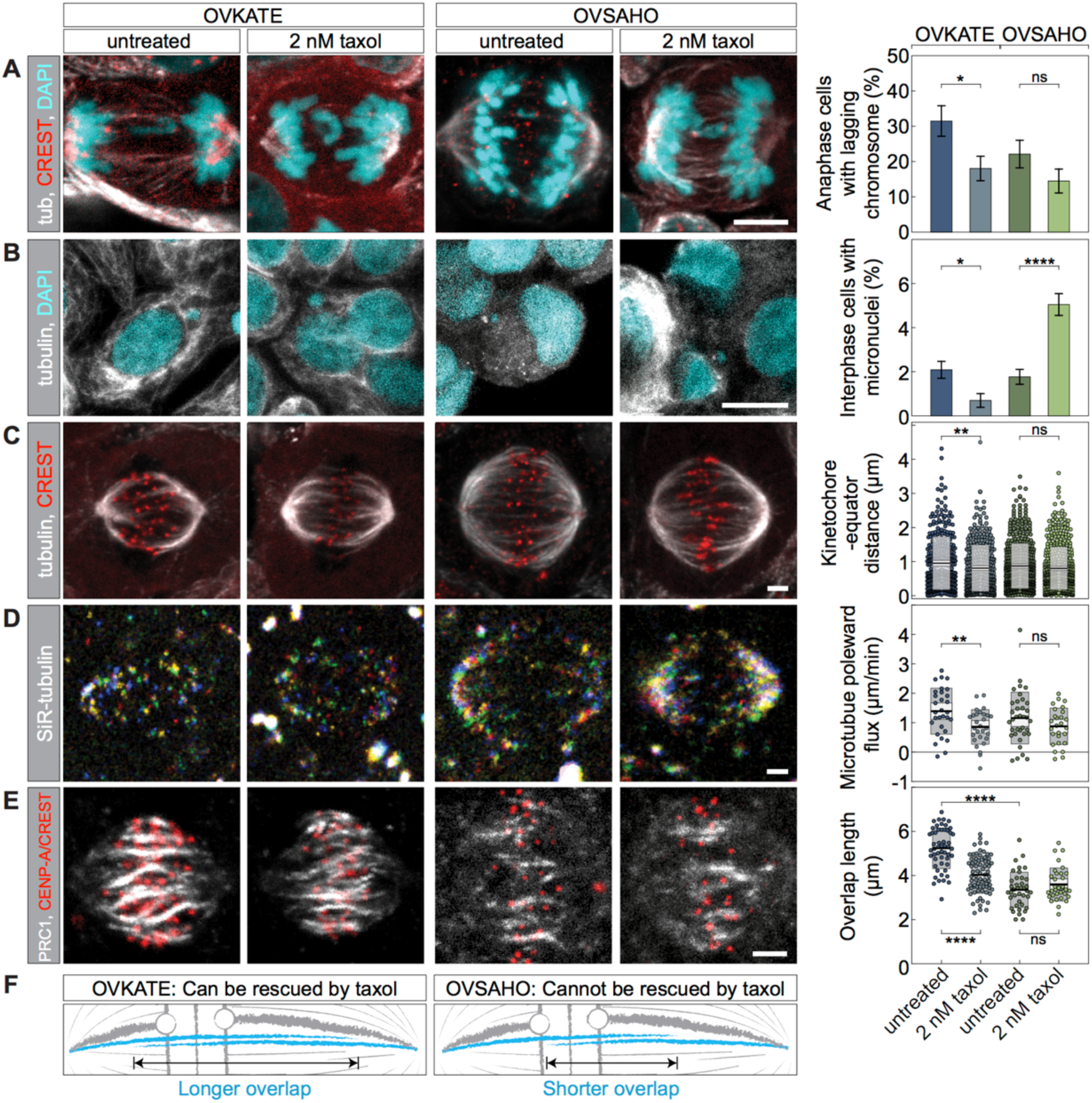
Modulation of chromosome missegregation rates by targeting microtubule poleward flux in ovarian cancer cells. **A,** OVKATE (left) and OVSAHO (right) cells immuno-stained for tubulin (grey), CREST (red), and dyed with DAPI (cyan). Untreated (left) and taxol-treated (right) cells are shown. Graph shows percentage of anaphase cells with lagging chromosome, color-coded for treatments. Number of analyzed cells for treatments from left to right: 100, 103, 96, 88. Scale bars: 5 µm. **B,** OVKATE (left) and OVSAHO (right) cells immuno-stained for tubulin (grey) and DAPI (cyan). Untreated (left) and taxol-treated (right) cells are shown. Graph shows percentage of interphase cells with micronuclei, color-coded for treatments. Number of analyzed cells for treatments from left to right: 1390, 716, 1527, 1961. Scale bars: 10 µm. **C,** OVKATE (left) and OVSAHO (right) cells immuno-stained for tubulin (grey) and CREST (red). Untreated (left) and taxol-treated (right) cells are shown. Graph shows kinetochore-equator distance, color-coded for treatments. Number of distances between sister kinetochore pairs and the equator measured for treatments from left to right: 337, 365, 507, 490. Scale bars: 2 µm. **D,** OVKATE (left) and OVSAHO (right) cells stained with SiR-tubulin (grey). Untreated (left) and taxol-treated (right) cells are shown. Temporal projection of SiR-tubulin speckles. Graph shows microtubule poleward flux rates, color-coded for treatments. Number of SiR-tubulin speckles tracked for treatments from left to right: 30, 28, 36, 27. Scale bars: 2 µm. **E,** OVKATE (left) and OVSAHO (right) cells immuno-stained for PRC1 (grey) and CENP-A in OVKATE (red) or CREST in OVSAHO (red). Untreated (left) and taxol-treated (right) cells are shown. Graph shows overlap length, color-coded for treatments. Number of PRC1-labelled bundles measured for treatments from left to right: 53, 74, 39, 33. Scale bars: 2 µm. **F,** Schematic illustrates that taxol treatment can reduce increased microtubule poleward flux and its adverse effects when overlap lengths are extended. Maximum intensity z-projection (A–E). Statistical tests: two-proportions z-test (A,B), Mann-Whitney test (C), t-test (D,E). p-value as in **Fig. 1**.

To assess whether the difference in the taxol effect on mitotic fidelity between the two cancer cell lines can be explained by our model, we tested the model’s predictions for these cases. The model suggests that taxol affects chromosome alignment, microtubule poleward flux, and antiparallel microtubule overlaps in OVKATE, but to a lesser extent or not at all in OVSAHO cells. Indeed, we found that taxol treatment improved chromosome alignment in OVKATE cells, whereas the improvement was smaller and not significant in OVSAHO cells (**Fig. 4C**; Supplementary Fig. S4D). Similarly, taxol decreased the microtubule poleward flux velocity in OVKATE cells, while the change was again smaller and not significant in OVSAHO cells (**Fig. 4D**; Supplementary Movie 3). Finally, we found that the origin of the differential effect of taxol treatment on microtubule poleward flux lies in the differential effect on overlap length, as overlaps were shortened by taxol in OVKATE cells, but not in OVSAHO cells (**Fig. 4E**; Supplementary Fig. S4E). Furthermore, the overlaps in untreated OVSAHO cells were shorter compared to those in OVKATE cells (**Fig. 4E**), implying that the initial overlap lengths may serve as an indicator of whether chromosome misalignment and missegregation can be rescued by reducing microtubule poleward flux.

Collectively, our results show that by modifying the length of antiparallel overlaps, which serve as a platform for motor proteins that drive microtubule poleward flux, chromosome missegregation rates can be altered in cancer cells. We propose that the difference in the effect of taxol between the two ovarian cancer cell lines is due to the fact that OVKATE cells possess lengthy antiparallel overlaps that can be shortened by taxol, leading to a slower microtubule poleward flux, rescuing chromosome alignment and maintaining segregation fidelity. Conversely, OVSAHO cells have short overlaps that remain unchanged after taxol treatment, lacking downstream mitotic effects and rescue. Thus, the inherent variations in spindle architecture, especially in microtubule overlaps, among cancer cell lines determine the efficacy of rescuing mitotic errors through the modulation of poleward flux (**Fig. 4F**).

## Discussion

In this paper, we show that overlap length-dependent microtubule poleward flux dictates chromosome segregation error and micronuclei formation frequencies via its effect on chromosome alignment (**Fig. 5**). Specifically, in two systems characterized by excessively long overlaps of antiparallel microtubules, KIF18A-depleted RPE1 cells and the ovarian cancer OVKATE cells, lagging chromosomes and micronuclei were rescued by treatments that shortened the overlaps and slowed down the poleward flux. We propose that when the overlap zones of antiparallel microtubules are shorter, the poleward flux is slower as a result of shorter overlaps between the antiparallel segments of bridging and k-fibers, which serve as a platform for motor proteins that slide these fibers apart and generate poleward flux of k-fibers (Risteski et al., 2022). We show that slower flux of k-fibers results in better chromosome alignment at the metaphase plate because k-fibers that flux slowly also grow slowly at the plus end, leading to slow kinetochore movements and thus less extensive oscillations around the spindle midplane, producing a tight metaphase plate. We suggest that this mechanism works together with the flux-driven centering (Risteski et al., 2022), where slower k-fiber flux promotes chromosome alignment because it implies that both k-fibers slip against the fast-fluxing bridging fibers. This allows the longer k-fiber to win in a tug-of-war against the shorter one and to move the chromosome towards the spindle midplane. As the microtubule poleward flux is controlled not only by the antiparallel overlaps, but also at the spindle pole, the finding that the pole-localized proteins WDR62 and katanin promote synchronous chromosome movement in anaphase (Guerreiro et al., 2021) highlights the role of the poleward flux in controlling mitotic fidelity.

**Fig. 5.**
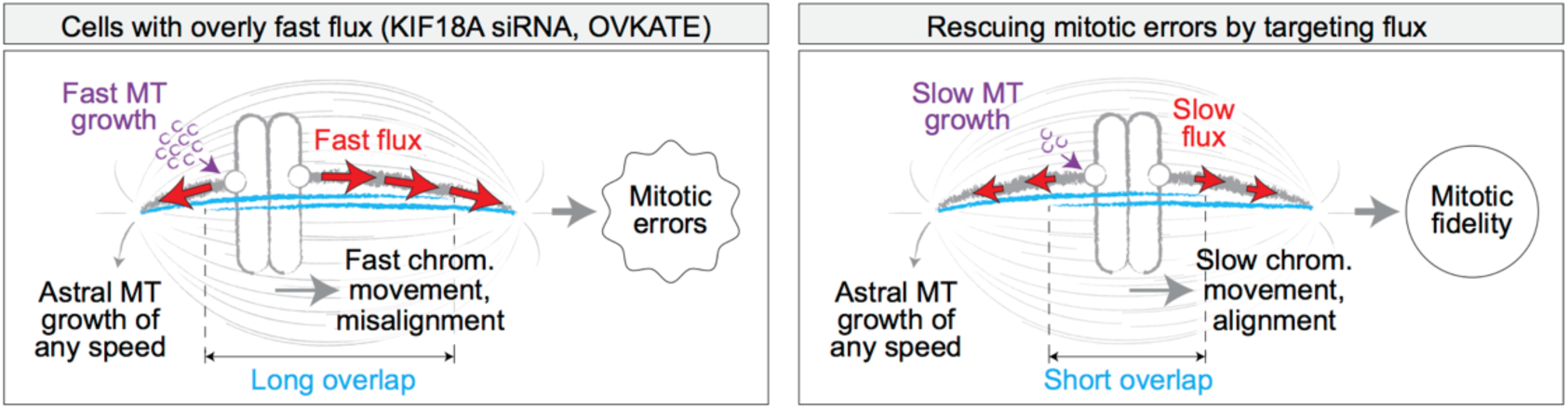
Enhancing mitotic fidelity by fine-tuning chromosome alignment through microtubule flux. In systems characterized by compromised mitotic fidelity, chromosome segregation errors may be mitigated through targeting elevated microtubule poleward flux caused by overly extended antiparallel microtubules. By shortening these overlap regions, where flux-driving forces are generated, excessive flux and, in turn, kinetochore microtubule growth are slowed down, ultimately leading to improved chromosome alignment on the spindle and fewer lagging chromosomes.

Previous studies have demonstrated that CIN in colorectal and high-grade serous ovarian cancer cells can be decreased by attenuating the increased microtubule dynamics in these cells by using a low-dose taxol treatment (Ertych et al., 2014; Tamura et al., 2020). Interestingly, in our KIF18A-depleted cells, a low-dose taxol also decreased CIN, but did not affect the overall microtubule dynamics, inferred from the growth rates of astral microtubules on metaphase spindles. Yet, taxol treatment decreased the growth rate of k-fibers at the plus end and their poleward flux velocities. Thus, we propose that a low-dose taxol treatment reduces mitotic errors by affecting the dynamics and flux of k-fibers rather than by changing the general dynamics of microtubules in systems characterized by excessively fast k-fiber flux, such as KIF18A-depleted cells.

Overexpression of KIF18A was detected in whole-genome duplicated tumors (Quinton et al., 2021), which could help tumor proliferation by avoiding micronuclei formation via dampened chromosome movements during metaphase. Cancer cell lines with dampened chromosome movements within the metaphase plate, e.g. HeLa – cervical adenocarcinoma (Ma et al., 2010) and U2OS – osteosarcoma (Ganem et al., 2005; Steblyanko et al., 2020) exhibit reduced microtubule poleward flux when compared to non-transformed RPE-1 – retinal pigment epithelium cells (Dudka et al., 2018; Risteski et al., 2022). This is in agreement with our model, where slower flux results in better chromosome alignment. In spite of a tight metaphase plate in these cancer cells, lagging chromosomes may appear as a consequence of inefficient error correction of kinetochore-microtubule attachments due to attenuated kinetochore oscillations (Iemura et al., 2021). However, as most cancer cell lines are transcriptionally distinct from their primary cancer types (Warren et al., 2021), it would be interesting to study the relationship between microtubule flux, chromosome alignment, and mitotic errors in patient tumor samples across various cancer types.

KIF18A promotes normal cell growth and viability as generation of micronuclei due to the lack of KIF18A hampers proliferation through a p53-dependent mechanism (Czechanski et al., 2015; Fonseca et al., 2019; Gomes et al., 2022; Sepaniac et al., 2021). In agreement with these studies, we found that majority of lagging chromosomes which arise due to chromosome alignment defects are able to integrate into the main nucleus, thus decreasing the frequency of micronuclei occurrence with respect to the occurrence of lagging chromosomes. In instances where micronuclei do form, the probability of micronuclei rupture may be a detrimental factor for cell proliferation. As the micronuclear envelope stability depends on chromosome length and gene density (Mammel et al., 2022), it would be interesting to test whether these micronuclei show bias towards certain chromosomes (Klaasen et al., 2022). Micronuclei resulting from chromosome misalignment form more stable nuclear envelopes than those resulting from defective kinetochore-microtubule attachments (Sepaniac et al., 2021). Unlike in normal tissues, micronuclear envelope integrity associated with insufficient KIF18A is compromised in tumors (Sepaniac et al., 2021). These factors underlying micronuclear integrity might influence cell proliferation following the modification of poleward flux.

By targeting microtubule poleward flux, chromosome alignment is affected independently of kinetochore attachment status (Fonseca et al., 2019; Risteski et al., 2022), and chromosome missegregation rates can be reduced in cells that have increased overlap lengths and microtubule poleward flux velocities. This suggests that microtubule poleward flux could be used as a target to mitigate or aggravate mitotic errors in cancers that show wider metaphase plates and longer spindles, where the latter serves as an indication of extended overlaps and increased microtubule poleward flux. Specifically, by increasing already elevated chromosome misalignment via targeting microtubule poleward flux, e.g. by affecting KIF18A, we speculate that it may be possible to increase the rate of chromosome missegregation beyond the tolerable threshold and promote cancer cell death (Cosper et al., 2022; Janssen et al., 2009; Kops et al., 2004). With the recent development of small-molecule inhibitors targeting KIF18A and their encouraging preclinical efficacy observed in certain human cancer models (Gliech et al., 2024; Payton et al., 2023), our study suggests that the misalignment of chromosomes together with elongation of antiparallel overlaps and spindles in cancer cells compared to their non-cancer counterparts, may signal vulnerability to KIF18A inhibition. These biophysical markers could serve as a basis for developing synthetic lethality strategies that include KIF18A inhibition, aiming to selectively eliminate cells exhibiting elevated microtubule poleward flux.

## Authors’ Contributions

**P. Risteski:** Conceptualization, formal analysis, investigation, visualization, methodology, writing–original draft, writing–review and editing. **J. Martinčić:** Conceptualization, formal analysis, investigation, visualization, methodology, writing– original draft. **E. Tintor:** Formal analysis, writing–original draft. **M. Jagrić:** Formal analysis, investigation, visualization, methodology. **I.M. Tolić:** Conceptualization, supervision, funding acquisition, writing–original draft, writing–review and editing.

## Acknowledgments

The authors thank Alexey Khodjakov (New York State Department of Health), Patrick Meraldi (University of Geneva), Marin Barišić (Danish Cancer Institute), and Dragomira Majhen (Ruđer Bošković Institute) for the cell lines, Anamaria Brozović (Ruđer Bošković Institute) for the chemicals, and Ivana Šarić (Ruđer Bošković Institute) for the drawings. The Tolić lab is funded by the European Research Council (ERC Synergy Grant, GA Number 855158), the Croatian Science Foundation (HRZZ) through Swiss-Croatian Bilateral Projects (project IPCH-2022-10-9344), and projects co-financed by the Croatian Government and the European Union through the European Regional Development Fund—the Competitiveness and Cohesion Operational Programme: IPSted (Grant KK.01.1.1.04.0057) and QuantiXLie Center of Excellence (Grant KK.01.1.1.01.0004).

## Methods

### Cell culture

The hTERT-RPE-1 cell line with stable expression of CENP-A-GFP and centrin1-GFP was a gift from Alexey Khodjakov (Wadsworth Center, New York State Department of Health, Albany, NY, USA). The hTERT-RPE-1 cell line expressing EB3-GFP and H2B-mCherry was a gift from Patrick Meraldi (University of Geneva, Geneva, Switzerland). Unlabeled human rhabdomyosarcoma and lung adenocarcinoma cell lines, RD and A549, respectively, were a gift from Dragomira Majhen (Ruđer Bošković Institute, Zagreb, Croatia). Unlabeled human colorectal adenocarcinoma cell line, DLD-1, was a gift from Marin Barišić (Danish Cancer Institute, Copenhagen, Denmark). Unlabeled human high-grade serous ovarian cancer cell lines, OVKATE and OVSAHO, were purchased from Tebu-Bio. RPE-1, RD, A549, and DLD-1 cells were cultured in Dulbecco’s Modified Eagle Medium (with 4.5 g/L d-glucose, stable glutamine, and sodium pyruvate; Capricorn Scientific), supplemented with 10% Fetal Bovine Serum (Sigma-Aldrich), 100 IU/mL penicillin, and 100 mg/mL streptomycin (Lonza). OVKATE and OVSAHO cell lines were cultured in RPMI 1640 medium (Sigma Aldrich) with the same supplements. Additionally, RPE-1 cells expressing EB3-GFP and H2B-mCherry were treated with 50 µg/mL of geneticin (Life Technologies). All cells were incubated at 37°C in a Galaxy 170 R humidified incubator (Eppendorf) with a 5% CO2 atmosphere. Routine mycoplasma testing was performed by visually inspecting DAPI staining under the microscope. Passages were limited to a maximum of 8–10 weeks (∼10 passages). 2 h prior to live cell imaging or fixation, cells were subjected to a low dose of paclitaxel (taxol; National Cancer Institute), specifically 2 nM (Crowley et al., 2022; Tamura et al., 2020).

### Cell transfection

The transfection of RPE-1 cell lines was carried out using the lipofection method. A day prior to siRNA transfection, 120,000 cells were seeded on 35-mm glass-bottom dishes with a glass thickness of 0.17 mm (ibidi GmbH). The siRNA constructs were diluted in Opti-MEM medium (Gibco), and transfection was performed with Lipofectamine RNAiMAX Reagent (Invitrogen) following the manufacturer’s protocol. The constructs and their final concentrations used were as follows: 100 nM KIF18A siRNA (4390825; Ambion), 100 nM KIF4A siRNA (sc-60888; Santa Cruz Biotechnology), 20 nM HAUS8 siRNA (L-031247-01-0005; Dharmacon), and 100 nM NuMA siRNA (sc-43978; Santa Cruz Biotechnology). After a 4-h incubation with the transfection mixture, the medium was replaced with regular cell culture medium. All experiments involving siRNA-treated cells were conducted 24 h after transfection, except for HAUS8, where silencing was carried out for 48 h. The effectiveness of siRNA silencing was previously validated with immunostaining (Risteski et al., 2022).

### Cell fixation and immunostaining

In experiments aimed at determining anaphase errors and kinetochores-to-equator distances, cells were fixed in pre-warmed 4% paraformaldehyde for 10 min. For assessing PRC1-labeled overlap length, cells were fixed with ice-cold methanol for 1 min. Following fixation, cells were permeabilized for 15 min in 0.5% Triton X-100 in PBS. Subsequently, cells underwent blocking with 1% NGS in PBS for 1 h and were then incubated with primary antibodies overnight at 4°C. Primary antibodies for PRC1, CREST, CENP-A, and tubulin were diluted at concentrations of 1:100, 1:300, 1:300, and 1:500, respectively, in 1% NGS in PBS. After the primary antibody incubation, cells were incubated with fluorescence-conjugated secondary antibodies at room temperature for 1 h. Secondary antibodies for PRC1, CREST, CENP-A and tubulin staining were prepared at dilutions of 1:250, 1:500, 1:500, and 1:1000, respectively, in 2% NGS in PBS. Following each step, cells were washed three times in PBS for 5 min. The primary antibodies used were mouse anti-PRC1 (sc-376983; Santa Cruz Biotechnology), rabbit anti-CENP-A (ab62242; Abcam), rat anti-alpha tubulin (MA1-80017, Invitrogen), and human anti-centromere protein antibody (Antibodies Incorporated). The secondary antibodies used were donkey anti-mouse IgG-Alexa Fluor 488 (Abcam), donkey anti-rabbit IgG-Alexa Fluor 488 (Abcam), donkey anti-mouse IgG-Alexa Fluor 594 (Abcam), donkey anti-mouse IgG-Alexa Fluor 647 (Abcam), donkey anti-rat IgG-Alexa Fluor 594 (Abcam), donkey anti-rat IgG-Alexa Fluor 647 (Abcam), and goat anti-human IgG H&L DyLight 594 (Abcam). Additionally, DAPI (1 μg/mL) was added and incubated for 15 min at room temperature to visualize chromosomes.

### Imaging of fixed cells

Immunostained cells, for determination of PRC1-labeled overlap length, were subjected to imaging using the Bruker Opterra Multipoint Scanning Confocal Microscope (Bruker Nano Surfaces) and the Zeiss LSM 800 confocal microscope (Zeiss). The Opterra Multipoint Scanning Confocal Microscope (Bruker Nano Surfaces) was mounted on a Nikon Ti-E inverted microscope equipped with a Nikon CFI Plan Apo VC 100x/1.4 numerical aperture oil objective (Nikon). The system was controlled with the Prairie View Imaging Software (Bruker). Images were captured with an Evolve 512 Delta EMCCD Camera (Photometrics, Tucson, AZ, USA). A 2x relay lens was placed in front of the camera, which resulted in the xy-pixel size in the image of 83 nm. Opterra Dichroic and Barrier Filter Set 405/488/561/640 was used to separate the excitation light from the emitted fluorescence with the following emission filters: BL HC 525/30, BL HC 600/37 and BL HC 673/11 (Semrock). The imaging was done on 5 central z-planes with 0.5-µm spacing. The Zeiss LSM 800 confocal microscope, with an Airyscan Superresolution Module and Axio Observer.Z1 inverted stand (Carl Zeiss GmbH, Germany), was also used for imaging tubulin-immunostained cells and observing anaphase chromosome errors. Only bipolar spindles were imaged, and they were imaged using multiple z-planes to encompass the entire anaphase spindle with all chromosomes. For excitation of blue, green, red and far-red fluorescence, 405/488/561/640 laser lines were used, respectively. Images were captured using the LSM 800 camera with Plan-Apochromat 63x/1.40 Oil DIC M27 objective (Carl Zeiss GmbH) or Plan-Apochromat 40x/0.95 Corr M27 objective. The acquisition of interphase cells for micronuclei detection was conducted on fixed cells using the Zeiss LSM 800 confocal microscope with 40x/0.95 Corr M27 objective and Tiles recording mode on multiple z-planes to encompass the entire cell volume. The system was operated with ZEN blue 3.4 system software (Carl Zeiss GmbH).

### Speckle microscopy

Cells cultured on glass-bottom dishes were treated with 1 nM SiR-tubulin dye (Spirochrome AG). Following a 15-min staining, confocal live imaging was conducted using a Dragonfly spinning disk confocal microscope system (Andor Technology) equipped with a 63x/1.49 HC PL APO oil objective (Leica) and Sona 4.2P scientific complementary metal oxide semiconductor camera (Andor Technology), or an Expert Line easy3D STED microscope system (Abberior Instruments) with a 60x/1.2 UPLSAPO 60XW water objective (Olympus) and avalanche photodiode detector. Fusion software and Imspector software were utilized for image acquisition, respectively. Throughout the imaging process, cells were maintained at a temperature of 37°C and a 5% CO2 atmosphere within a heating chamber (Okolab). For live imaging of RPE-1 cells expressing CENP-A-GFP and centrin1-GFP, and dyed with SiR-tubulin, the Dragonfly microscope system utilized 488-nm and 640-nm laser lines for the excitation of GFP and SiR, respectively. Similarly, the Expert Line microscope system

utilized 485-nm and 640-nm laser lines for the excitation of GFP and SiR, respectively. To visualize SiR-tubulin speckles, images were acquired with 80% laser power and an exposure time of 1 s. Image acquisition was performed on one or two focal planes every 5 or 10 s. To visualize the dynamics of speckles in the OVKATE and OVSAHO cell lines, SPY555-DNA dye (Spirochrome AG) was used to stain cell DNA at a final concentration of 100 nM, 2 h before imaging. To prevent dye efflux, a broad-spectrum efflux pump inhibitor, verapamil (Spirochrome AG), was introduced at a final concentration of 0.5 μM along with the DNA dye. Once the dye permeated the cells and metaphase spindles became discernible, 1 nM SiR-tubulin dye was introduced to visualize the dynamics of tubulin speckles. The imaging of speckles and DNA was conducted using 561-nm and 640-nm laser lines on the Dragonfly microscope system.

### Imaging of EB3 comets

The velocity of EB3 comets on astral microtubules was assessed in RPE-1 cells stably expressing EB3-GFP and H2B-mCherry. Metaphase cells were imaged every 2 s in a single z-plane for 1 min using the Dragonfly spinning disk confocal microscope system (Andor Technology). Image acquisition was operated with Fusion software (Andor Technology). Throughout the imaging process, cells were maintained at a temperature of 37°C and a 5% CO2 atmosphere within a heating chamber (Okolab). For excitation, 488-nm and 561-nm laser lines were used for visualization of GFP, and mCherry, respectively. A single central spindle z-plane, where both centrosomes were visible, was imaged with both laser lines every 2 s.

### Image and data analysis

Image analysis and measurements were performed in Fiji/ImageJ (National Institutes of Health). Quantification and data analysis were performed in R (R Foundation for Statistical Computing) and MATLAB (MathWorks). Figures and schemes were assembled in Adobe Illustrator CC (Adobe Systems). Statistical analysis was performed using Student’s t-test, Mann-Whitney test and two-proportions z-test.

Anaphase chromosome errors were quantified by examining fixed RPE-1 anaphase cells with stable expression of CENP-A-GFP and centrin1-GFP, which were subsequently stained for tubulin and DNA. In the case of unlabeled cancer cells, fixation was followed by staining for tubulin, DNA, and CREST. Analysis was restricted to bipolar spindles, involving the inspection of multiple z-planes of spindles during anaphase. A chromosome was categorized as “lagging” if it trailed the dividing chromosome mass by at least 2 μm. If there was no gap in the DNA signal between two chromosome arms, these chromosomes were classified as chromosome bridges and excluded from the calculation. Only spindles not in contact with the telophase contractile ring were included in the analysis.

To quantify interphase micronuclei, images of fixed cells were examined in the DNA channel. The *Multi-point tool* was utilized to mark each cell nucleus and determine the total cell count in the image. All slices in the Z-stack were inspected for micronuclei, and the occurrence of at least one micronucleus in a cell was documented. The tubulin channel was additionally inspected to confirm whether two or more micronuclei were associated with a single cell or multiple cells.

To measure kinetochore alignment on the spindle, we used the *Multi-point tool* to track the positions of sister kinetochore pairs. For RPE-1 cells, the measurements were obtained from live imaging, whereas the position of kinetochores in cancer cell lines was determined post-immunolabeling with the CREST antibody. The equatorial plane was defined as the line passing through two points positioned between the outermost pairs of kinetochores on opposite sides of the spindle. Chromosome alignment was then calculated as the distance between the midpoint of sister kinetochore pairs and the equatorial plane. Note that sister kinetochore pairs located near the centrosomes were excluded from consideration to prevent the tracking of kinetochores that are not in a bi-oriented configuration.

To assess the antiparallel overlap length in cells immunostained for PRC1, a 5-pixel-thick segmented line was used to trace the pole-to-pole contour within individual PRC1-labeled overlap regions. The pole-to-pole tracking was performed on single z-planes, and the mean value of the cytoplasmic intensity was subtracted from the obtained intensity profiles. The overlap length of individual PRC1-labeled regions was defined as the width of the intensity peak at the central part of the contour in SciDavis (Free Software Foundation Inc.). The width measurement was taken at the base of the PRC1 intensity peak, where the PRC1 signal roughly equals the mean value of the PRC1 signal along the contour on either side of the peak.

To assess microtubule poleward flux velocity using the speckle microscopy assay, we considered tubulin speckles that could be tracked for at least 30 s within the spindle. For each tubulin speckle position in RPE-1 cells, corresponding CENP-A and centrin positions, indicative of sister kinetochores and spindle poles, respectively, were tracked using the *Multi-point tool*. Speckles originating at a proximal kinetochore and associated with its proximal pole were categorized as part of the k-fiber, while those starting between sister kinetochores or proximal to the distal pole and passing through sister kinetochores were classified as part of the bridging fiber. In unlabeled cancer cell lines where kinetochores were not visible during live imaging, only the positions of speckles and spindle poles were tracked over time. Speckle-pole velocity was determined by fitting linear regression to the distances between the SiR-tubulin speckle and the associated spindle pole during the first 30 s of its trajectory. From the corresponding speckle trajectories, kinetochore positions were used to calculate kinetochore velocity. This was achieved by fitting a linear regression to the distances between kinetochores and the pole over the 30-s timeframe. The kinetochore was categorized as “leading” if it moved towards its associated pole and “trailing” if it moved away from its associated pole throughout the 30-s speckle trajectory. To ensure that kinetochores were actively leading or trailing, without being in a pausing state, a threshold was set. Kinetochores were selected based on a movement criterion within 10 s of at least 300 nm towards or away from the pole for leading and trailing kinetochores, respectively. The speckle-kinetochore velocity was then determined by fitting linear regression to the distances between the speckle and the associated kinetochore over the 30-s period.

EB3 velocity of astral microtubules was assessed in RPE-1 cells expressing EB3-GFP and H2B-mCherry. The positioning of chromosomes within the spindle was imaged to identify metaphase spindles. The trajectories of EB3 comets on astral microtubules were acquired on a single z-plane using the *Multipoint tool*. Tracking of the comets and their corresponding spindle poles was conducted over time, starting from the initial appearance of the comet to the last frame where the comet remained clearly visible. The tracking spanned at least 4 time points (8 s), and the comet velocity was determined by fitting linear regression to the comet-pole distances over time. Kymographs for EB3 comets were generated using a 10-pixel thick *Straight Line*, drawn from pole to pole, and then applying the *KymographBuilder* plugin.

## Supplementary Material

**Supplementary Fig. S1.**
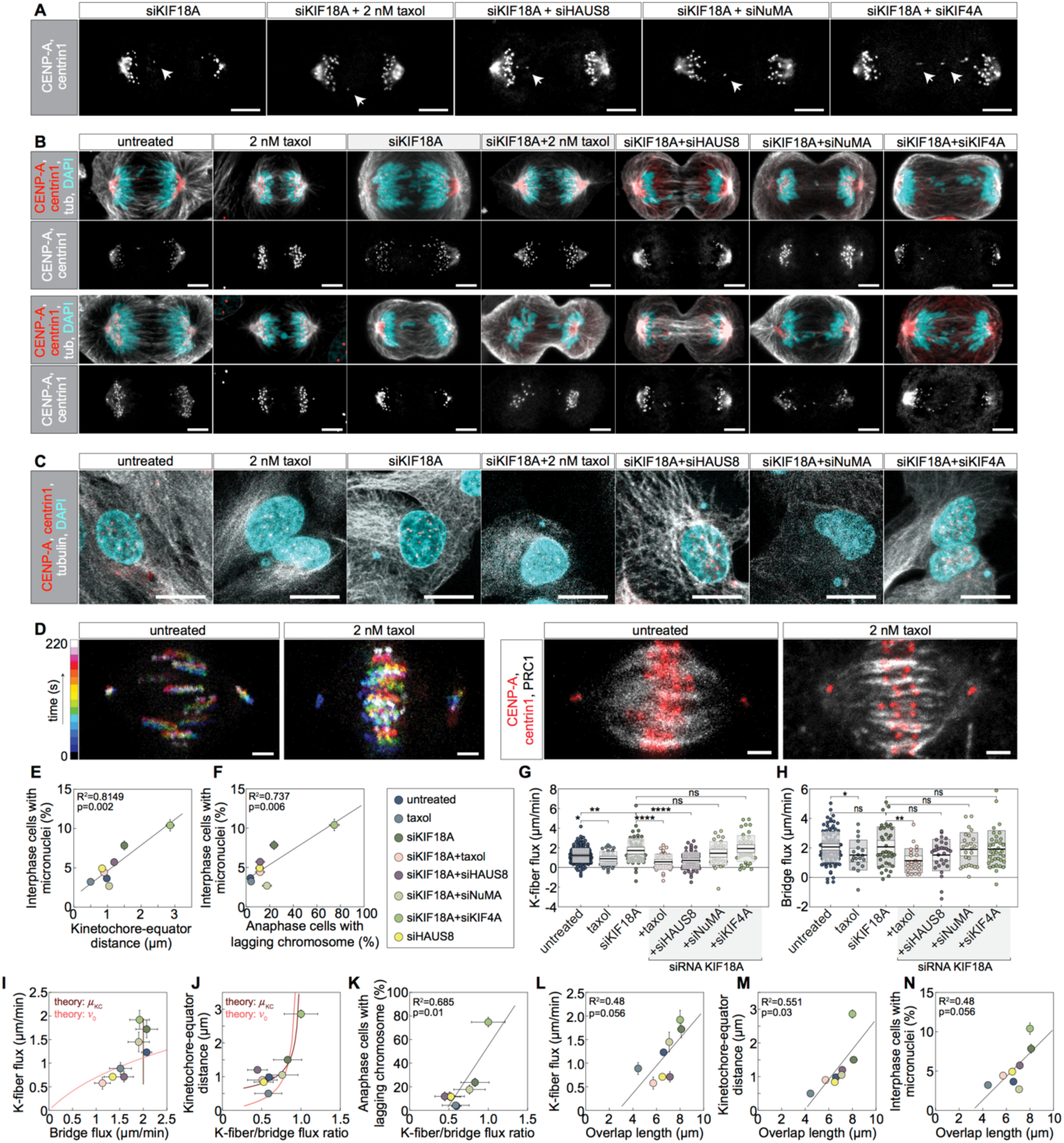
Microtubule flux parameters and overlap lengths in adverse effects of chromosome misalignment. **A,** RPE-1 cells from **Fig. 1A**. Individual CENP-A-GFP, centrin1-GFP channel of corresponding treatments is shown. **B,** Lagging chromosomes per corresponding treatment in RPE-1 cells with stable expression of CENP-A-GFP and centrin1-GFP (red), immuno-stained for tubulin-AF-594 (grey) and dyed with DAPI (cyan). Individual CENP-A-GFP, centrin1-GFP channel is shown below merged channels. Scale bars: 5 µm. **C,** Micronuclei in RPE-1 cells with stable expression of CENP-A-GFP and centrin1-GFP (grey), immuno-stained for tubulin-AF-594 (grey) and dyed with DAPI (cyan), per corresponding treatment. **D,** Temporal projection of live kinetochore and centrosome movements (left), and PRC1-labeled overlaps (right) per corresponding treatment in RPE-1 cells with stable expression of CENP-A-GFP and centrin1-GFP (red), immuno-stained for PRC1-AF-594 (grey). Scale bars: 2 µm. **E,** Percentage of interphase cells with micronuclei plotted against the kinetochore-equator distance, color-coded for treatments. **F,** Percentage of interphase cells with micronuclei plotted against the percentage of anaphase cells with lagging chromosomes, color-coded for treatments. **G,** Measurement of k-fiber flux rates per corresponding treatment. Number of analyzed k-fiber speckles for treatments from left to right: 164, 35, 52, 40, 60, 35, 43, 87. **H,** Measurement of bridging fiber flux rates per corresponding treatment. Number of analyzed bridging fiber speckles for treatments from left to right:101, 19, 37, 24, 30, 25, 36, 39. **I,** Graph showing dependence of k-fiber flux rates on bridging fiber flux rates. **J,** Graph showing dependence of the kinetochore-equator distance on k-fiber to bridging fiber flux ratio. **K,** Graph showing dependence of percentage of cells with anaphase lagging chromosomes on k-fiber to bridging fiber flux ratio. **L,** Graph showing dependence of k-fiber flux rate on overlap length. **M,** Kinetochore-equator distance plotted against the overlap length. **N,** Percentage of interphase cells with micronuclei plotted against the overlap length. Legend is at the bottom central part. Maximum intensity z-projection (A–C, D; right). Kinetochore-equator distances, overlap lengths, and flux for untreated, siKIF18A, siKIF18A+siHAUS8, siKIF18A+siKIF4A, and siHAUS8 were replotted from (Risteski et al., 2022). Statistical tests: t-test (G, H). p-value as in **Fig. 1**.

**Supplementary Fig. S2.**
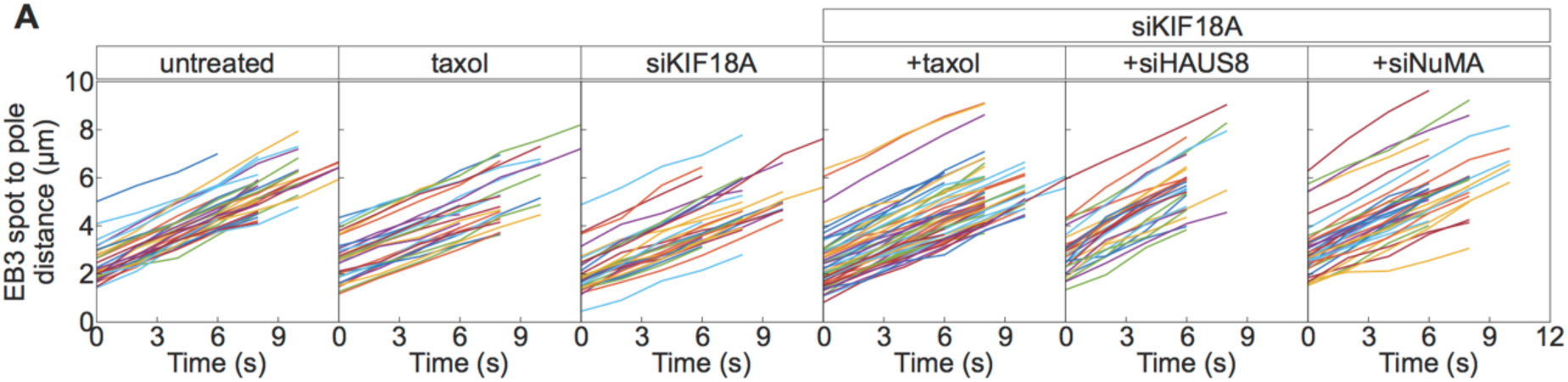
Assessment of astral microtubule plus-end dynamics. **A,** EB3 spot to pole distances over time per treatment. EB3 spots on astral microtubules in metaphase were used; different colors represent individual EB3 tracks. The tracks were used to calculate EB3 spot velocity in **Fig. 2B**.

**Supplementary Fig. S3.**
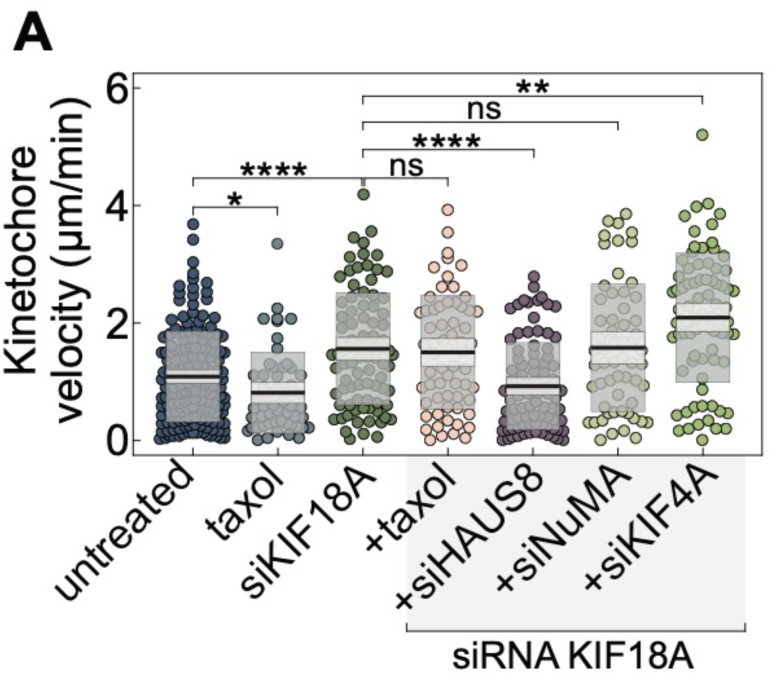
Assessment of kinetochore velocities. **A,** Kinetochore velocity on the spindle. Number of measured sister kinetochore velocities for treatments from left to right corresponds to number of tracked k-fiber speckles in **Supplementary Fig. S1G**. Statistical tests: t-test. p-value as in **Fig. 1**.

**Supplementary Fig. S4.**
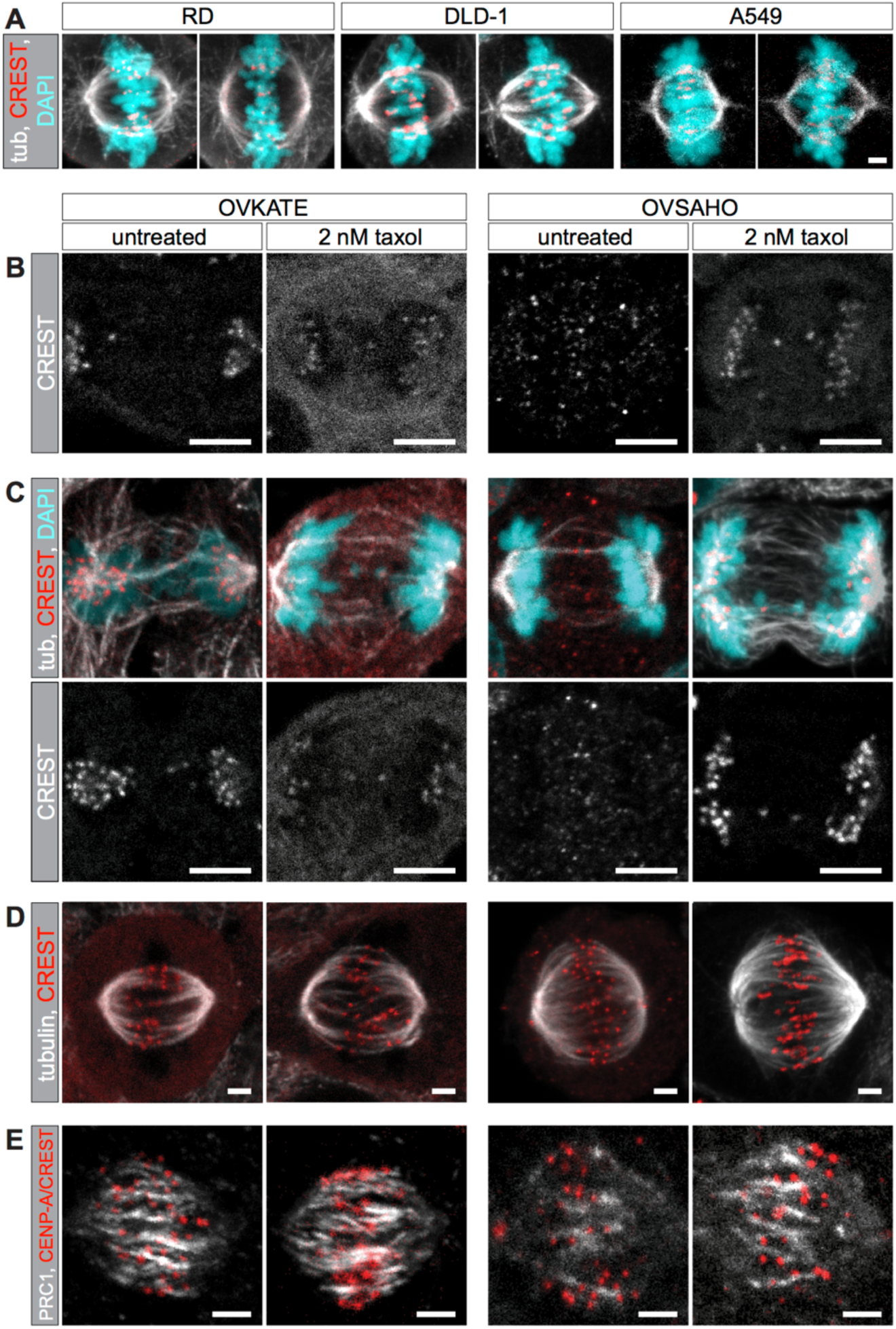
Assessment of overlap length-dependent chromosome misalignment in cancer cell lines. **A,** RD (left), DLD-1 (middle), and A549 (right) cell lines stained with tubulin (grey), CREST (red), and dyed with DAPI (cyan). **B,** OVKATE (left) and OVSAHO (right) cells from **Fig. 4A**. Individual CREST channels of untreated (left) and taxol-treated (right) cells are shown. Scale bars: 5 µm. **C,** Additional examples of cells as in **Fig. 4A**. Scale bars: 2 µm. **D,** Additional examples of cells as in **Fig. 4C**. Scale bars: 2 µm. **E,** Additional examples of cells as in **Fig. 4E**. Scale bars: 2 µm. Maximum intensity z-projection (A–E).

### Supplementary Movies 1–3

**Supplementary Movie 1. Metaphase kinetochore movement in cells with KIF18A-depleted background.** Montage of movies of RPE-1 cells stably expressing CENP-A-GFP (grey) and centrin1-GFP (grey) used for temporal color projection in **Fig. 1A**. Single z-plane. Time interval: 5 s. Scale bar: 2 µm.

**Supplementary Movie 2. EB3 plus end dynamics in cells with KIF18A-depleted background.** Montage of movies of RPE-1 cells stably expressing EB3-GFP (grey) and H2B-mCherry (cyan) used for temporal projection in **Fig. 2A**. Single z-plane. Time interval: 2 s. Scale bar: 2 µm.

**Supplementary Movie 3. Microtubule poleward flux in ovarian cancer cell lines.** Montage of movies of OVKATE and OVSAHO cells stained with SiR-tubulin (grey) and SPY555-DNA (cyan) used for temporal color projection in **Fig. 4D**. Untreated and taxol-treated cells are shown. Maximum intensity z-projection of 2 planes. Time interval: 5 s. Scale bar: 2 µm.

